# The Effect of MR Image Quality on Structural and Functional Brain Connectivity: The Maastricht Study

**DOI:** 10.1101/806075

**Authors:** Joost J.A. de Jong, Jacobus F.A. Jansen, Laura W.M. Vergoossen, Miranda T. Schram, Coen D.A. Stehouwer, Joachim E. Wildberger, David E.J. Linden, Walter H. Backes

## Abstract

In large population-based cohort studies, magnetic resonance imaging (MRI) is often used to study the structure and function of the brain. Advanced MRI techniques such as diffusion-tensor (dMRI) or resting-state functional MRI (rs-fMRI) can be used to study connections between distinct brain regions. However, brain connectivity measures are likely affected by biases introduced during MRI data acquisition and/or processing.

We identified three sources that may lead to bias, i.e. signal-to-noise ratio (SNR), head motion, and spatial mismatch between MRI-based anatomy and a brain atlas. After quantifying these sources, we determined the associations between the image quality metrics and brain connectivity measures derived from dMRI and rs-fMRI in 5,110 participants of the population-based Maastricht Study.

More head motion and low SNR were negatively associated with structural and functional brain connectivity, respectively, and these metrics substantially affected (>10%) associations of brain connectivity with age, sex and body mass index (BMI), whereas associations with diabetes status, educational level, history of cardiovascular disease, and white matter hyperintensities were less or not affected. In addition, age, sex, and BMI were associated with head motion, SNR, and atlas mismatch (all p < 0.001). Based on our results, we strongly advise that, in large population-based cohort neuroimaging studies, statistical analyses on structural and functional brain connectivity should adjust for potentially confounding effects of image quality.

**Highlights:**

- Low MR image quality compromises brain connectivity measures
- MR image quality is negatively associated with age, body mass index, and male sex
- Statistical analyses in large neuroimaging studies should account for image quality

## Introduction

### Neuroimaging in population studies

Population-based cohort studies are extremely relevant sources of fundamental research data, contribute to a better understanding of health effects of life styles and pathophysiology of diseases, and reveal key information on risk factors (Szklo, 1998). If structure and function of the brain are of interest, neuroimaging using magnetic resonance imaging (MRI) is often the preferred tool to incorporate into the study design. Neuroimaging can provide valuable structural and functional information on the brain, but the large amount of individuals in combination with the typical size of MRI data poses certain challenges in terms of data acquisition, storage, processing, and analysis (Smith and Nichols, 2018). Although there are several on-going large-scale neuroimaging population-based cohort studies, e.g. the Generation R Study (White et al., 2013), Rotterdam Scan Study (Ikram et al., 2015), UK Biobank (Miller et al., 2016), Human Connectome Project (Van Essen et al., 2013; Van Essen et al., 2012), The Rhineland Study (Breteler et al., 2014), and The Maastricht Study (Schram et al., 2014), each using different scanner hardware, study-specific scan protocols, and processing tools, there is no consensus on data acquisition and processing. In order to recognize potential biases introduced during the data acquisition and processing, it is important to be transparent about the quality of the MRI data itself and the way these data are processed.

### Brain connectivity

In the last two decades, advanced MRI techniques have been developed that allow mapping of the connectivity of the brain’s network. Two main techniques to do so are typically diffusion-weighted MRI (dMRI) and resting-state functional MRI (rs-fMRI). dMRI estimates the axonal orientations which are consecutively used to calculate white matter fiber tracts between brain regions, i.e. structural connectivity, using tractography algorithms (Basser et al., 1994; Jones et al., 1999). Rs-fMRI data are used to calculate functional connectivity as the correlation between temporal changes in blood-oxygen-level-dependent (BOLD) signal of spatially distinct brain regions (Biswal et al., 1995; Friston et al., 1993).

Recent research has indicated that not only in neurological and psychiatric disorders (van den Heuvel and Hulshoff Pol, 2010), but also in systemic conditions such as type-2 diabetes mellitus (T2DM), structural (Hoogenboom et al., 2014; Reijmer et al., 2013; van Bussel et al., 2016a) as well as functional brain connectivity (Supekar et al., 2008; van Bussel et al., 2016b; Wang et al., 2006; Zhang et al., 2010) are altered compared to healthy controls.

### Objectives

Both dMRI and rs-fMRI rely on echo planar imaging (EPI) pulse sequences, which are known to exhibit various forms of image degradation or artefacts to a greater extent than structural MR imaging (Jezzard and Balaban, 1995). Therefore, the determination of brain connectivity outcome measures from these images might be affected, for instance due to signal-to-noise ratio (SNR) limitations (DeDora et al., 2016; Dikaios et al., 2014), head motion (Baum et al., 2018; Satterthwaite et al., 2012; Van Dijk et al., 2012), and magnetic field inhomogeneities leading to geometric distortions, which in turn can result in misalignment (spatial mismatch) between dMRI and rs-fMRI data and brain atlases (Despotovic et al., 2015; Wang and Yushkevich, 2012).

To quantify these potential sources of bias in dMRI and rs-fMRI data of The Maastricht Study (Schram et al., 2014), we implemented a quality assessment procedure within the structural and functional brain connectivity processing pipeline. The aim of the current study was to investigate whether structural and functional connectivity outcome measures are affected by these image quality metrics and, if they are, analyse how to account for this bias. We therefore posed the following research questions: 1) Are the image quality metrics noise, head motion and atlas mismatch related to measures of structural and functional brain connectivity?; 2) Does image quality affect associations between brain connectivity and typical demographic variables of interest, i.e. age, sex, body mass index, diabetes status, educational level, history of cardiovascular disease, and white matter hyperintensities?; and 3) Which of these demographic variables are associated with low image quality?

## Methods

### Study population

We used data from the Maastricht Study, an observational population-based cohort study. The rationale and methodology have previously been described (28). In brief, the study focuses on the etiology, pathophysiology, complications, and comorbidities of type 2 diabetes and is characterized by an extensive phenotyping approach. Eligible for participation were all individuals aged between 40 and 75 years and living in the southern part of the Netherlands. Participants were recruited through mass media campaigns, the municipal registries, and the regional Diabetes Patient Registry via mailings. Recruitment was stratified according to known type 2 diabetes status, with an oversampling of individuals with type 2 diabetes for reasons of efficiency. Structural, diffusion and resting-state functional MRI measurements were implemented from December 2013 onward until February 2017 and were completely available in 5,261 of 5,547 participants. Processing of the dMRI or rsfMRI failed in 71 participants and in the remaining 5,190 participants, complete data on covariates were available in 5,110 participants (a flow-chart is given in **Supplemental Figure S1** in the appendix). The study has been approved by the institutional medical ethics committee (NL31329.068.10) and the Minister of Health, Welfare and Sports of the Netherlands (permit 131088-105234-PG). All participants gave written informed consent.

### MRI data acquisition and retrieval

For each participant, MRI data were acquired on a 3T clinical magnetic resonance scanner (MAGNETOM Prisma^fit^, Siemens Healthineers GmbH, Munich, Germany) located at a dedicated scanning facility (Scannexus, Maastricht, The Netherlands) using a head/neck coil with 64 elements for parallel imaging. The MRI protocol included a three-dimensional (3D) T1-weighted (T1w) magnetization prepared rapid acquisition gradient echo (MPRAGE) sequence (repetition time/inversion time/echo time (TR/TI/TE) 2,300/900/2.98ms, 176 slices, 256 × 240 matrix size, 1.0 mm cubic reconstructed voxel size); a fluid-attenuated inversion recovery (FLAIR) sequence (TR/TI/TE 5,000/1,800/394 ms, 176 slices, 512 × 512 matrix size, 0.49 × 0.49 × 1.0 mm reconstructed voxel size); a resting-state functional MRI (rs-fMRI) using a task-free T2*-weighted blood oxygen level-dependent (BOLD) sequence (TR/TE 2,000/29 ms, flip angle 90°, 32 slices (interleaved acquisition order), 104 × 104 matrix size, 2.0 × 2.0 × 4.0 mm reconstructed voxel size, 195 dynamic volumes); and a diffusion-tensor MRI (dMRI) using a diffusion sensitized echo-planar imaging (EPI) sequence (TR/TE 6,100/57 ms, 65 slices, 100 × 100 matrix size, 64 diffusion sensitizing gradient directions (b=1,200 s/mm2), 2.0 mm cubic reconstructed voxel size) with additionally three minimally-diffusion-weighted images (b=0 s/mm2).

Contraindications for MRI assessments were the presence of a cardiac pacemaker or implantable cardioverter defibrillator, neurostimulator, nondetachable insulin pump, metallic vascular clips or stents in the head, cochlear implant, metal-containing intrauterine device, metal splinters or shrapnel, dentures with magnetic clip, an inside bracket, pregnancy, epilepsy, and claustrophobia.

### Segmentation of brain tissue

T1w and FLAIR data were analysed by use of an ISO13485:2012–certified, automated method (which included visual inspection)(de Boer et al., 2009; Vrooman et al., 2007). T1w data were segmented into gray matter, white matter, white matter hyperintesities (WMH), and CSF volumes (1 voxel = 1.00 mm3 = 0.001 mL) (de Boer et al., 2009). Intracranial volume was calculated as the sum of gray matter, white matter (including WMH volume), and CSF volumes.

### dMRI and rs-fMRI data pre-processing

dMRI as well as rs-fMRI data were first anonymized and converted from DICOM to NIfTI format using Chris Rorden’s dcm2nii tool (version 2MAY2016 64bit BSD License) for further processing.

Pre-processing of the dMRI data was mainly performed with ExploreDTI v4.8.6 (PROVIDI lab, Image Sciences Institute, Utrecht, The Netherlands) (Leemans et al., 2009), and included eddy current and head motion correction (Farrell et al., 2007; Leemans and Jones, 2009), followed by constrained spherical deconvolution (CSD)-based deterministic whole-brain tractography (Tax et al., 2014) to obtain white matter fiber tracts. Next, the automated anatomical labelling (AAL) atlas (Rolls et al., 2015), consisting of 94 (sub)cortical brain regions in the cerebrum, was linearly coregistered to the dMRI data using FLIRT (Jenkinson and Smith, 2001) in FMRIB Software Library (FSL) 5.0.10 (FMRIB Analysis Group, University of Oxford, Oxford, U.K.). Lastly, for each pair of brain regions with two or more tracts running between them, the connection strength was determined as tract volume (number of voxels visited by a tract multiplied by the voxel size) relative to ICV, resulting in a symmetric 94×94 connectivity matrix, i.e. the participant’s structural connectome (SC), where each row and column represent a brain region and each element represents the relative tract volume between two regions.

Pre-processing of the rs-fMRI data was performed using a combination of tools in FSL 5.0.10 and Statistical Parametric Mapping (SPM) 12 (The Wellcome Trust College London, London, U.K.), and included magnetization stabilization followed by correction for field inhomogeneities (Zhang et al., 2001), slice-timing, and head motion (Soares et al., 2016). Next, rs-fMRI data were spatially and temporally filtered to increase signal-to-noise ratio (SNR) and remove possible respiratory and signal drift effects to focus on the spontaneous low-frequency fluctuations (Biswal et al., 1995). Lastly, the AAL atlas and individual-specific T1w including WM and CSF masks were linearly coregistered to the rs-fMRI data using FSL’s FLIRT (Jenkinson and Smith, 2001), and average time-series for each brain region as well as for the CSF and WM were calculated from the per-voxel time-series in each region. For each pair of brain regions, the connection strength was defined as the Pearson’s correlation coefficient calculated using linear regression of the averaged time-series of each region, corrected for motion (three translational and three rotational parameters) as well as the CSF and WM signal, resulting in the participant’s functional connectome. Negative correlations, which are considered not representing any meaningful connections, were set to zero (Smith et al., 2011).

In both the structural as well as the functional connectome, self-self connections, i.e. the diagonal elements, were set to zero. A complete overview of the structural and functional connectivity processing pipeline, including a description of the hardware and software, is given in the appendix (**S2**), and a schematic overview is shown in **Figure 1**.

**Figure 1.**
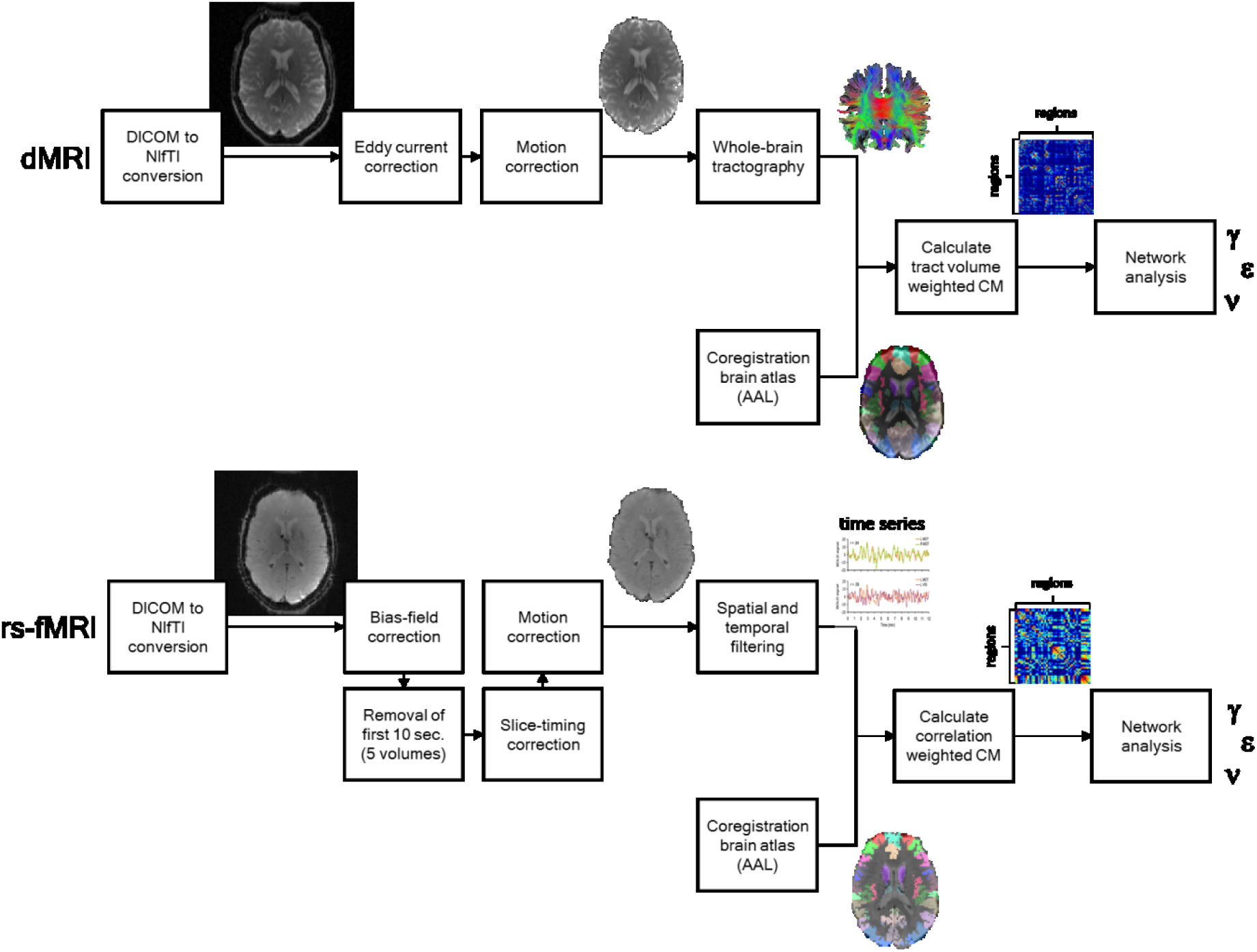
Schematic overview of the complete processing pipelines of the dMRI (top) and rs-fMRI (bottom) data to analyse structural and functional network connectivity, respectively.

### Brain network connectivity analysis using graph theory

From here on, the approach to calculate the structural and functional connectivity using graph theory was similar. First, one structural and one functional group-averaged connectome were calculated from all individual structural (n = 5,226) and functional (n = 5,231) connectomes, respectively. For the structural group-averaged connectome, the individual connectomes were used in binarized form (relative tract volume > 0), whereas for the functional group-averaged connectome the individual connectomes were used as such. To minimize the effect of spurious connections, both group-averaged connectomes were proportionally thresholded to a default sparsity of 0.80, meaning that only the connections that were present in at least 80% of the participants were taken into account in the individual structural and functional connectivity analyses. A schematic representation of the structural and functional group-averaged connectomes at sparsity 0.80 is shown in **Figure 2**.

**Figure 2.**
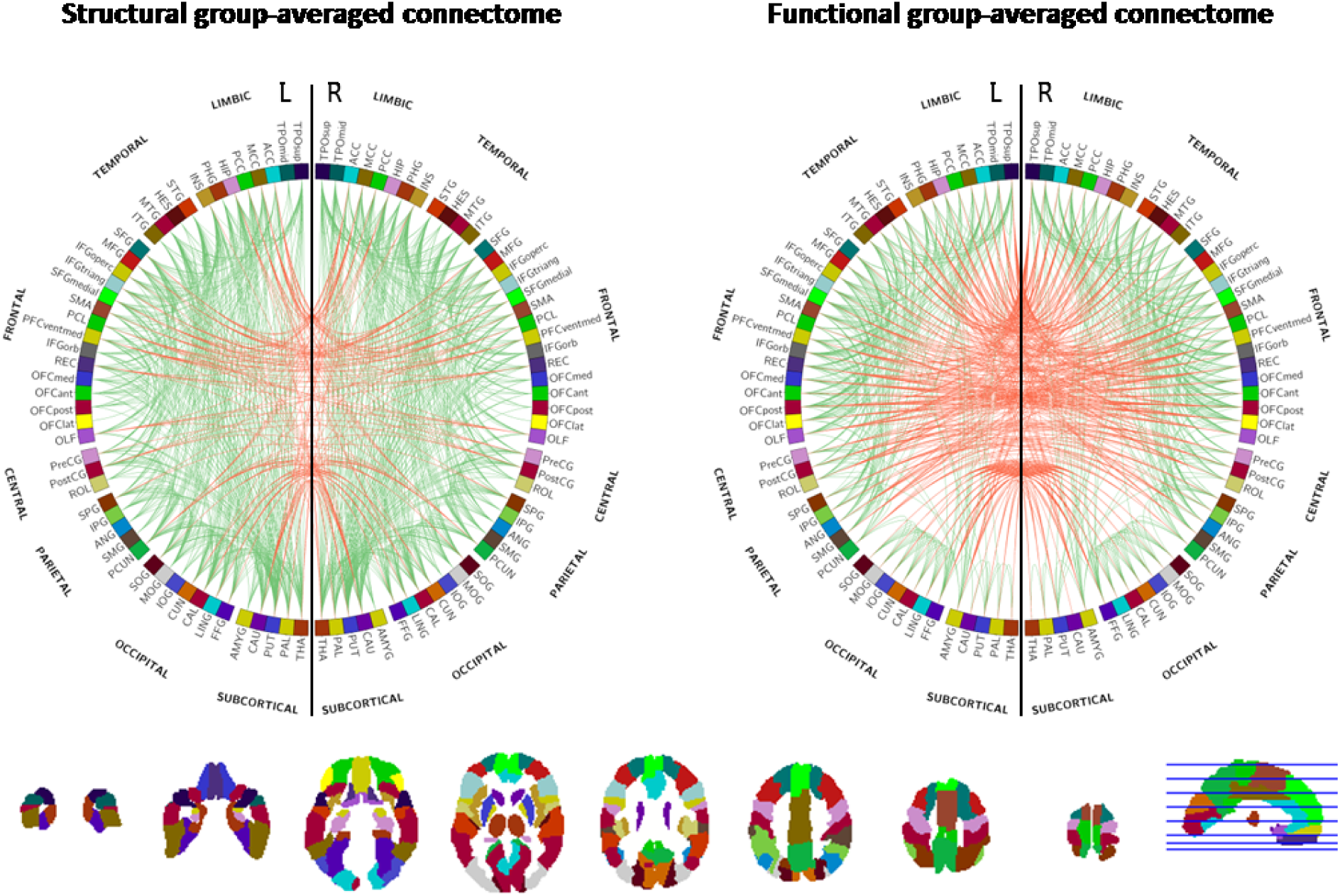
Structural (left) and functional (right) group-averaged connectomes at a sparsity of 0.80 showing the connections (# = 874) between brain regions that were subsequently used in the structural and functional network connectivity analyses, respectively. (Red: interhemispheric connections, green: intrahemispheric connections). Brain region labels and colours are according to the automated anatomical labeling (AAL2) atlas which is shown underneath.

Before thresholding the individual connectomes with the group-averaged connectome (Vasa et al., 2017), the participant’s structural and functional overall connectivity were calculated as the mean from all weights in the SC and FC, respectively (van den Heuvel et al., 2017). Subsequently, each participant’s connectome was masked by the group-averaged connectome, resulting in a weighted, undirected network with a sparsity close to the sparsity of the group-averaged connectome.

From each masked individual connectome, the following theoretical network connectivity measures were calculated using graph theory: average node degree (ν), a basic global network measure that can be interpreted as the “wiring cost” of the network (Rubinov and Sporns, 2010); normalized clustering coefficient (γ), a global measure of network segregation (Onnela et al., 2005; Rubinov and Sporns, 2010); and normalized global efficiency (ε_global_), a global measure of network integration (Latora and Marchiori, 2001; Maslov and Sneppen, 2002). The clustering coefficient and global efficiency were normalized to values calculated from 100 randomly generated networks of the same size, sparsity and binary degree as the individual network (Maslov and Sneppen, 2002; Rubinov and Sporns, 2010). All connectivity analyses were performed using the Brain Connectivity Toolbox (Rubinov and Sporns, 2010) in MATLAB Release 2016a (The Mathworks Inc., Natick, Massachusetts, U.S.).

To assess robustness of the connectivity measures over sparsity, the structural and functional group-averaged connectomes were additionally thresholded to sparsities ranging from 0.60 to 0.90 (step size 0.05) and from 0.10 to 0.90 (step size 0.10), respectively, and the connectivity measures were calculated at each of these sparsity values.

### Quality assessment

Uncertainty in brain connectivity measures was assessed using the following image quality metrics, each on a ‘lower is better’ scale: 1) inverse signal-to-noise ratio (iSNR) of the unprocessed images, 2) amount of head motion, and 3) spatial mismatch between the pre-processed dMRI or rs-fMRI data and the AAL brain atlas:

#### 1) Inverse signal-to-noise ratio

The iSNR [-] was calculated according to equation 4.1 (Association, 2008):

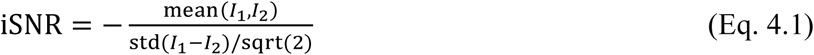

where *I_1_* and *I_2_* were two volumes that were acquired immediately after each other at b=0 s/mm^2^ at the end of the dMRI scan. For the rs-fMRI scan, *I_1_* and *I_2_*were the first two volumes that were acquired after removal of the first 10 seconds to account for magnetic stabilization, i.e. the 5^th^ and 6^th^ volume.

#### 2) Head motion

The amount of head motion was expressed as mean volume-to-volume translation, which was calculated from the translational parameters from the rigid body correction for head motion according to equation 4.2 (Van Dijk et al., 2012). In short, the translational head motion [mm] of a volume was computed as the root-mean-square of displacements in the sagittal, coronal, and transverse planes:

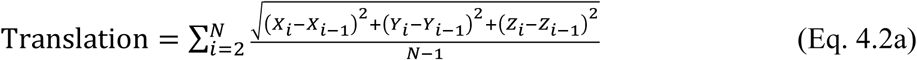

where *N* is the number of volumes in the dMRI or rs-fMRI data; and *X*, *Y*, and *Z* are the displacements of the *i*^th^ volume along the left-right, anterior-posterior, and longitudinal axis, respectively.

#### 3) Mismatch between brain atlas and pre-processed data

The spatial mismatch between the pre-processed dMRI or rs-fMRI data and the AAL brain atlas was quantified using 1-Dice’s similarity coefficient (Dice, 1945) according to equation 4.3:

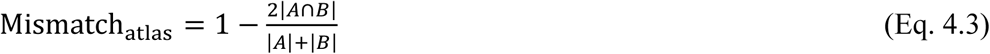

where *A* is the number of voxels in the brain mask of the dMRI or rs-fMRI data and *B* is the number of voxels in the brain mask of the AAL template. Mismatch varies between 0 and 1 representing no and complete mismatch, respectively.

### Demographic and clinical variables

Demographic and clinical data were collected as previously described (Schram et al., 2014). Variables of interest included age, sex, body mass index (BMI), educational level (‘Low’, ‘Middle’, or ‘High’) and history of cardiovascular disease (‘No’, or ‘Yes’). Based on their glucose metabolism status as determined according to the World Health Organization’s criteria by a 75-grams two-hour glucose tolerance test (OGTT) after an overnight fast (World Health Organization & International Diabetes Federation, 2006), participants were categorized into either ‘No diabetes’ (normal glucose metabolism), ‘Prediabetes’, ‘Type 2 diabetes’, or ‘Other type of diabetes’ (Schram et al., 2014).

### Statistics

Structural and functional brain connectivity measures were reported using the appropriate descriptive statistics, e.g. means and standard deviation in case of normally distributed data, median and 25^th^–75^th^ percentiles for non-normally distributed data, or percentages for categorical data.

Multiple linear regression was used to assess the relationship between quality metrics and structural and functional connectivity measures.

To study the effect of image quality on the association between brain connectivity and demographic and clinical variables, two linear regression models were used. In model 1, the connectivity measure was the independent variable and age, sex, BMI, diabetes status, educational level, history of CVD, and WMH volume were the dependent variables. In model 2, we additionally adjusted for the image quality metrics. Skewed variables (WMH volume) were log10-transformed. Significant regression coefficients that changed more than 10% were considered as relevant changes.

To ascertain which demographic and clinical variables age, sex, BMI, diabetes status, educational level, history of CVD, and WMH volume were associated with the quality metrics, linear regression was used. Skewed variables (WMH volume) were log10-transformed.

All statistical analyses used a level of significance of 0.05, and were performed in IBM SPSS Statistics for Windows, version 25 (IBM Corp., Armonk, NY, USA).

## Results

Demographic and clinical characteristics, brain connectivity estimates at a sparsity of 0.80, and dMRI and rs-fMRI image quality metrics in the participants that were included in this study (n = 5,110) are listed in **Table 1**.

**Table 1.** Characteristics of participants with successfully processed dMRI and rs-fMRI data (n=5,110).

| **Characteristic** |  |
| --- | --- |
| ***Demographic*** |  |
| Age [mean (SD), years] | 59.4 (8.7) |
| Sex [%] |  |
| Male | 50.6 |
| Female | 49.4 |
| Educational level [%]* |  |
| Low | 32.0 |
| Medium | 28.4 |
| High | 39.7 |
| ***Clinical*** |  |
| BMI [mean (SD), kg/m<sup>2</sup>] | 26.6 (4.2) |
| Diabetes status [%] |  |
| No diabetes | 64.1 |
| Prediabetes | 14.7 |
| Type 2 diabetes | 20.6 |
| Other type of diabetes | 0.6 |
| History of CVD [%]** |  |
| No | 87.4 |
| Yes | 12.6 |
| Relative WMH volume<br>[median (25<sup>th</sup>-75<sup>th</sup> percentile), % of ICV] | 0.016 (0.005-0.050) |
| ***Brain connectivity*** |  |
| Structural connectivity [mean (SD), -] |  |
| Overall | 5.3*10<sup>-3</sup> (0.6*10<sup>-3</sup>) |
| Average node degree, $\nu$ | 17.8 (0.4) |
| Clustering coefficient, $\gamma$ | 2.31 (0.08) |
| Global efficiency, $\epsilon_{\text{global}}$ | 0.84 (0.03) |
| Functional connectivity [mean (SD), -] |  |
| Overall | 0.32 (0.03) |
| Average node degree, $\nu$ | 16.7 (0.7) |
| Clustering coefficient, $\gamma$ | 3.25 (0.23) |
| Global efficiency, $\epsilon_{\text{global}}$ | 0.75 (0.02) |
| ***Image quality*** |  |
| **dMRI** |  |
| Signal-to-noise ratio [mean (SD), -] | 22 (6.5) |
| Head motion [mean (SD), mm] | 0.64 (0.16) |
| Atlas mismatch [mean (SD), -] | 0.087 (0.0092) |
| **Rs-fMRI** |  |
| Signal-to-noise ratio [mean (SD), -] | 39 (11) |
| Head motion [mean (SD), mm] | 0.13 (0.088) |
| Atlas mismatch [mean (SD), -] | 0.089 (0.0086) |

Abbreviations: SD: standard deviation; BMI: body mass index; CVD: cardiovascular disease; WMH: white matter hyperintensities; ICV: intracranial volume; dMRI: diffusion-tensor magnetic resonance imaging; Rs-fMRI: resting-state functional magnetic resonance imaging.
Missing data: \* Educational level (N=62); \*\* History of CVD (N=57).

**Figure 3** shows histograms of dMRI and rs-fMRI image quality metrics SNR, head motion and atlas mismatch.

**Figure 3.**
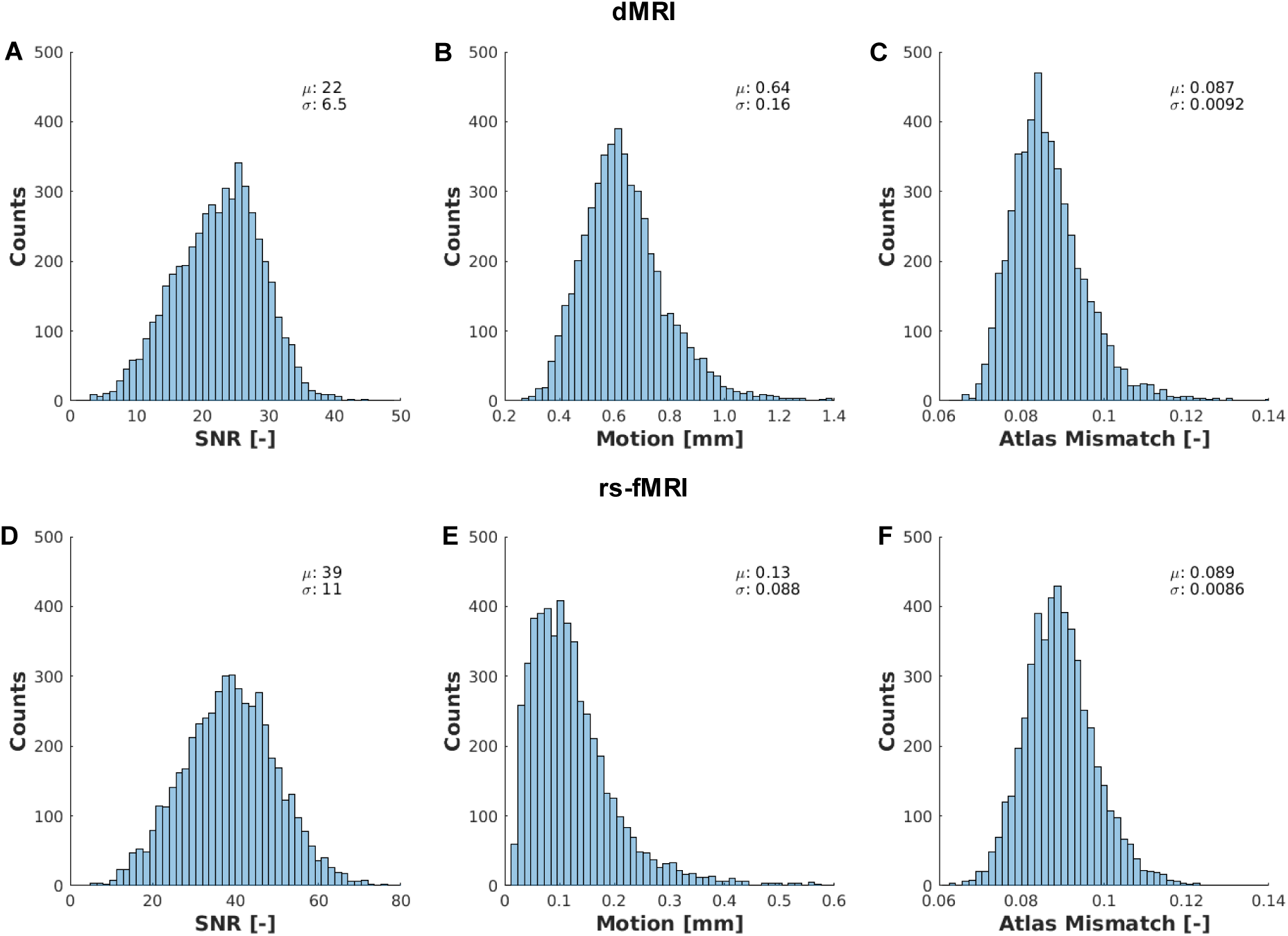
Histograms of dMRI (top) and rs-fMRI (bottom) image quality metrics signal-to-noise ratio (SNR) (left), amount of head motion (middle), and atlas mismatch (right).

In **Figure 4**, mean and 5^th^–95^th^ percentiles of structural and functional connectivity measures ν, γ and ε_global_ are plotted over the range of sparsities. Mean (standard deviation (SD)) of overall structural and functional connectivity (note that these are sparsity-independent) were 5.3*10^-3^ (0.6*10^-3^) and 0.32 (0.03), respectively.

**Figure 4.**
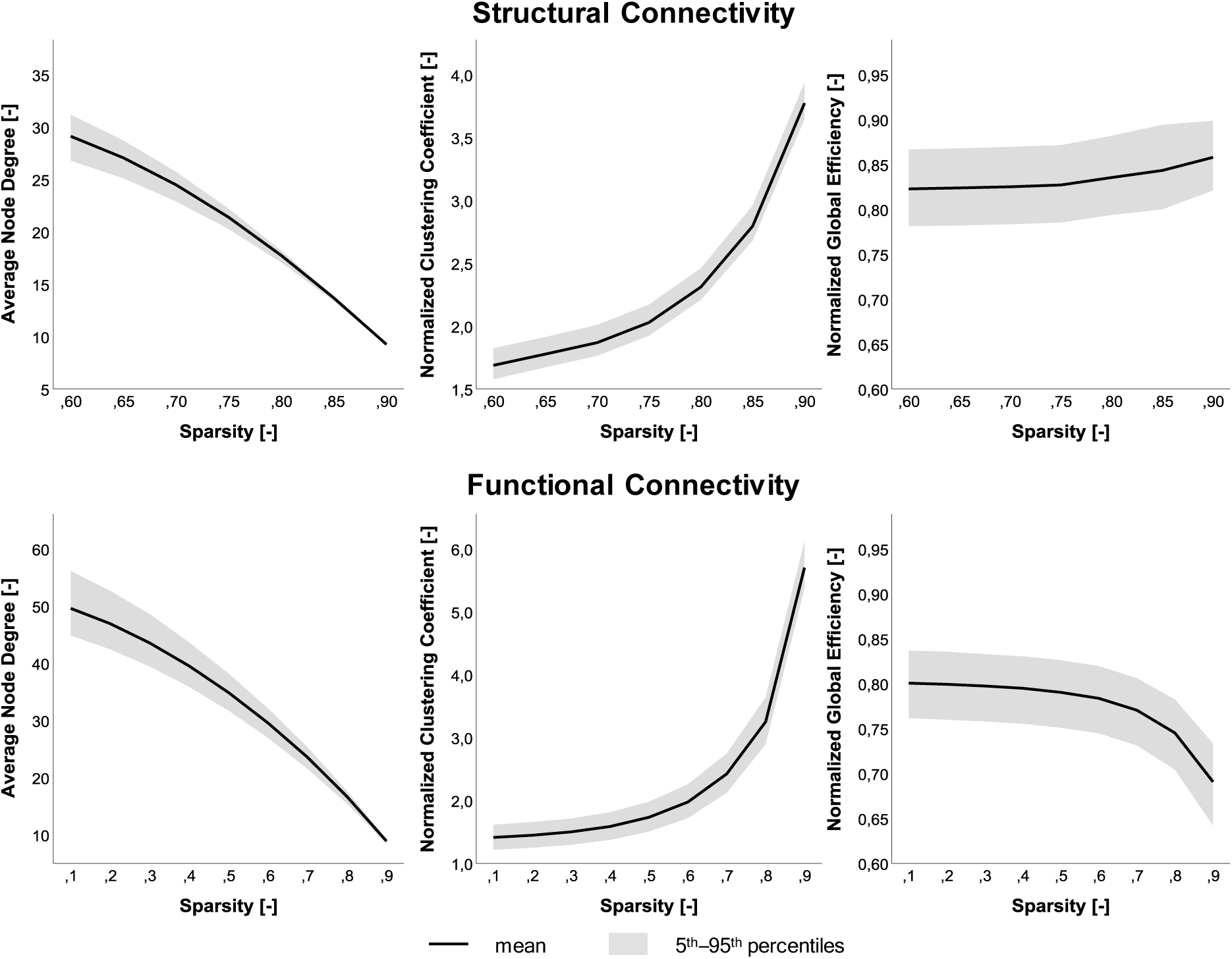
Mean and 5^th^–95^th^ percentiles of structural (top) and functional (bottom) connectivity measures ν (left), γ (middle), and ε_global_ (right) over the range of sparsities.

### Associations of connectivity measures with dMRI and rs-fMRI quality metrics

The diffusion MR image quality metrics iSNR, head motion and atlas mismatch were all related to structural connectivity measures overall SC, ν, and γ, with the strongest associations for head motion with standardized regression coefficients (β) ranging from –0.36 to 0.40 (all p<0.001), while atlas mismatch was most strongly related to ε_global_ (β = –0.15, p<0.001), as shown in **Figure 5**. A full overview of the associations between the diffusion and functional MR image quality metrics and the structural and functional connectivity measures, respectively, as well as the R^2^ for each model, is reported in the appendix (**Supplemental Table 3**).

**Figure 5.**
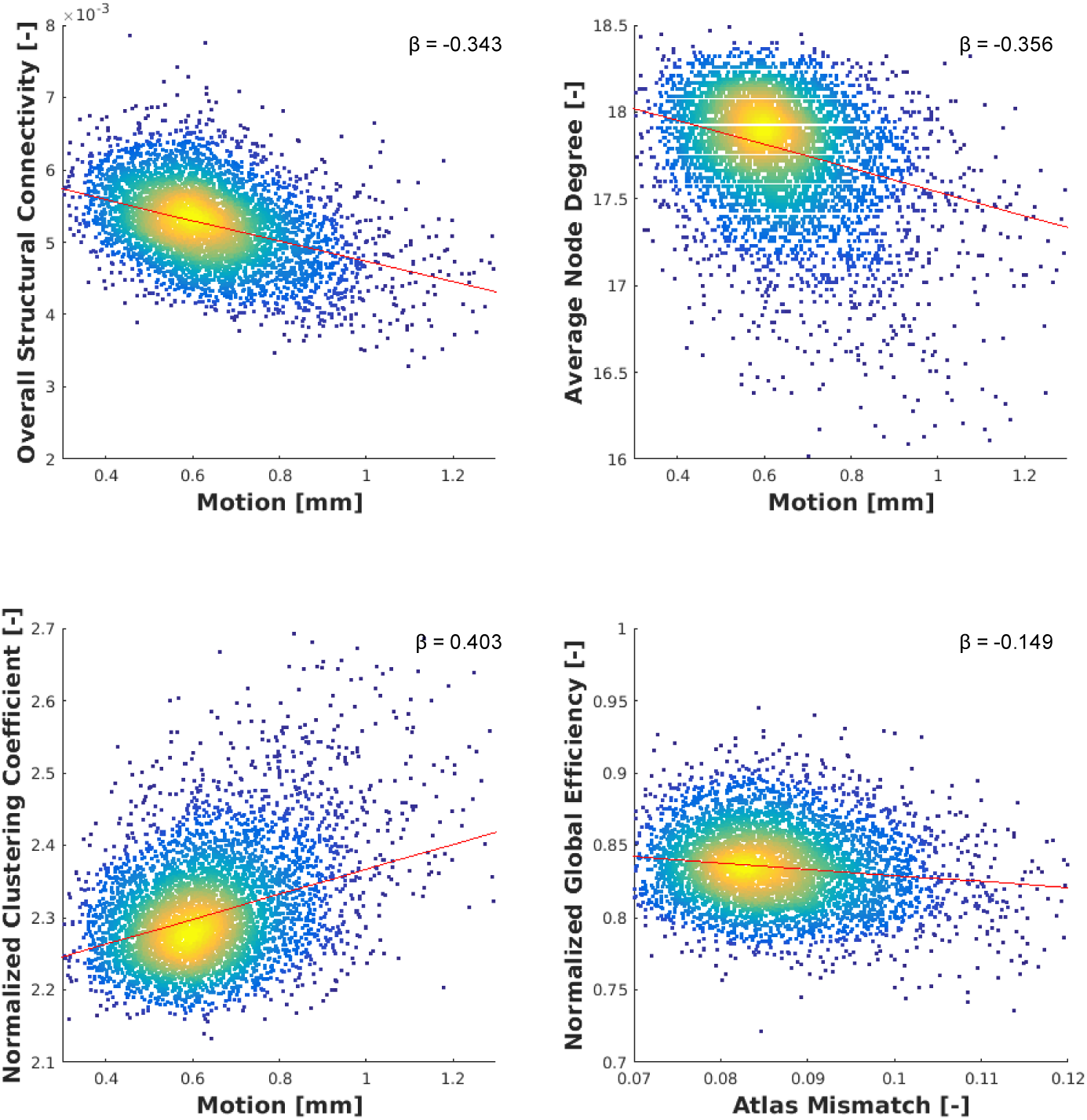
Density scatterplots of structural connectivity measures at sparsity 0.80 versus the quality metrics with the strongest association.

From the functional MR image quality metrics, iSNR was most strongly related to each of the functional connectivity measures overall FC, ν, and γ, with standardized regression coefficients (β) ranging from –0.22 to 0.15 (all p<0.001), except to ε_global_, for which head motion had the strongest association (β=0.16, p<0.001), as shown in **Figure 6**.

**Figure 6.**
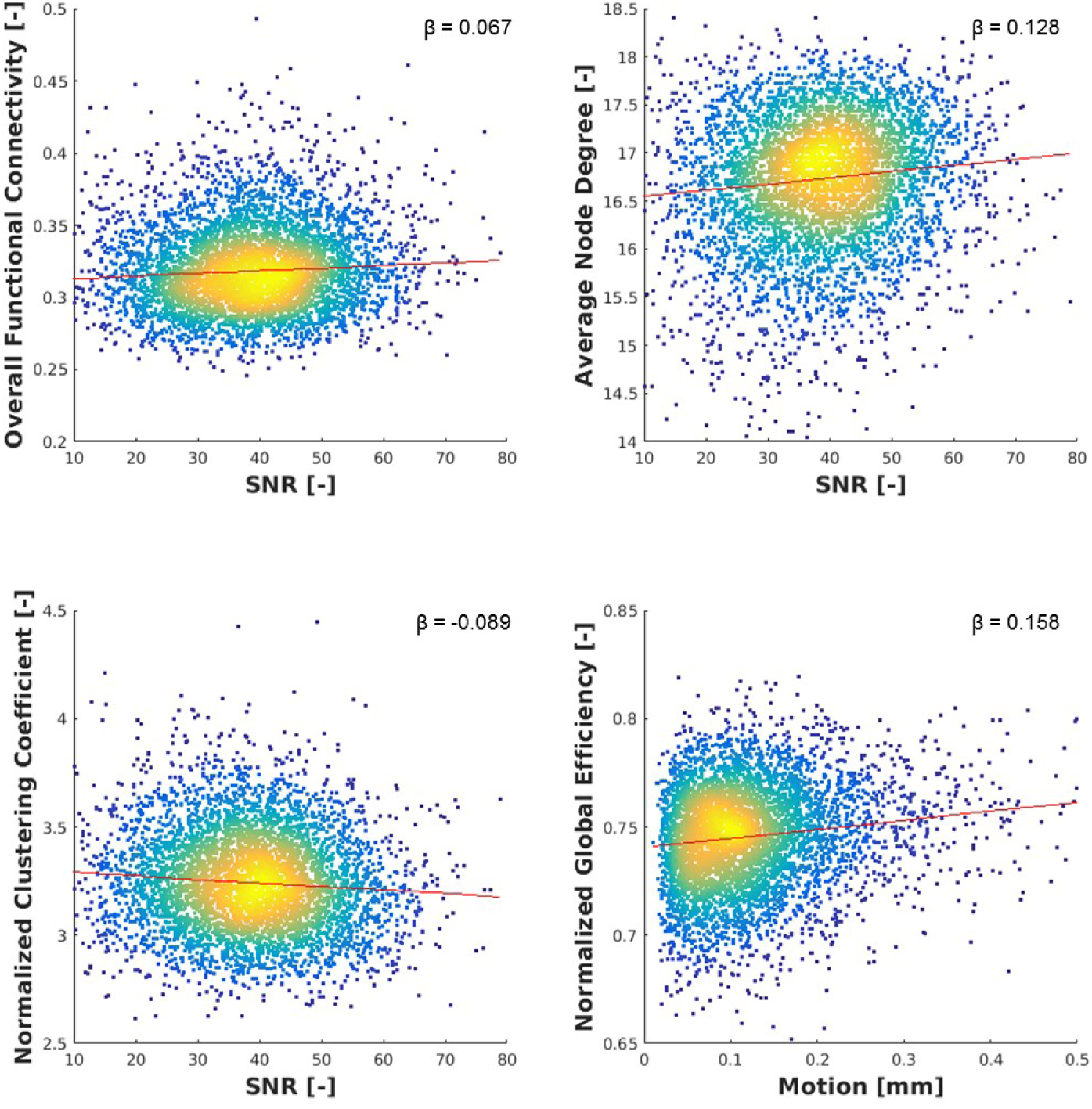
Density scatterplots of functional connectivity measures at sparsity 0.80 versus the quality metrics with the strongest association. Note: for intuitiveness, SNR (expressed as – iSNR) is plotted instead of iSNR.

The variability in the SC measures was consistently better explained by the quality metrics than variability in the FC measures, with R^2^-values ranging from 0.030 to 0.173 (3.0% to 17.3%) for the SC measures (see **Supplemental Table 4A**) and 0.006 to 0.032 (0.6% to 3.2%) for the FC measures (see **Supplemental Table 4B**).

### Effect of dMRI and rs-fMRI quality on connectivity associations

Standardized regression coefficients of the regression model between the structural or functional connectivity measures and the demographic/clinical variables, and the same model with additional adjustment for the image quality metrics are shown in **Table 2A** and **2B**.

**Table 2A.**
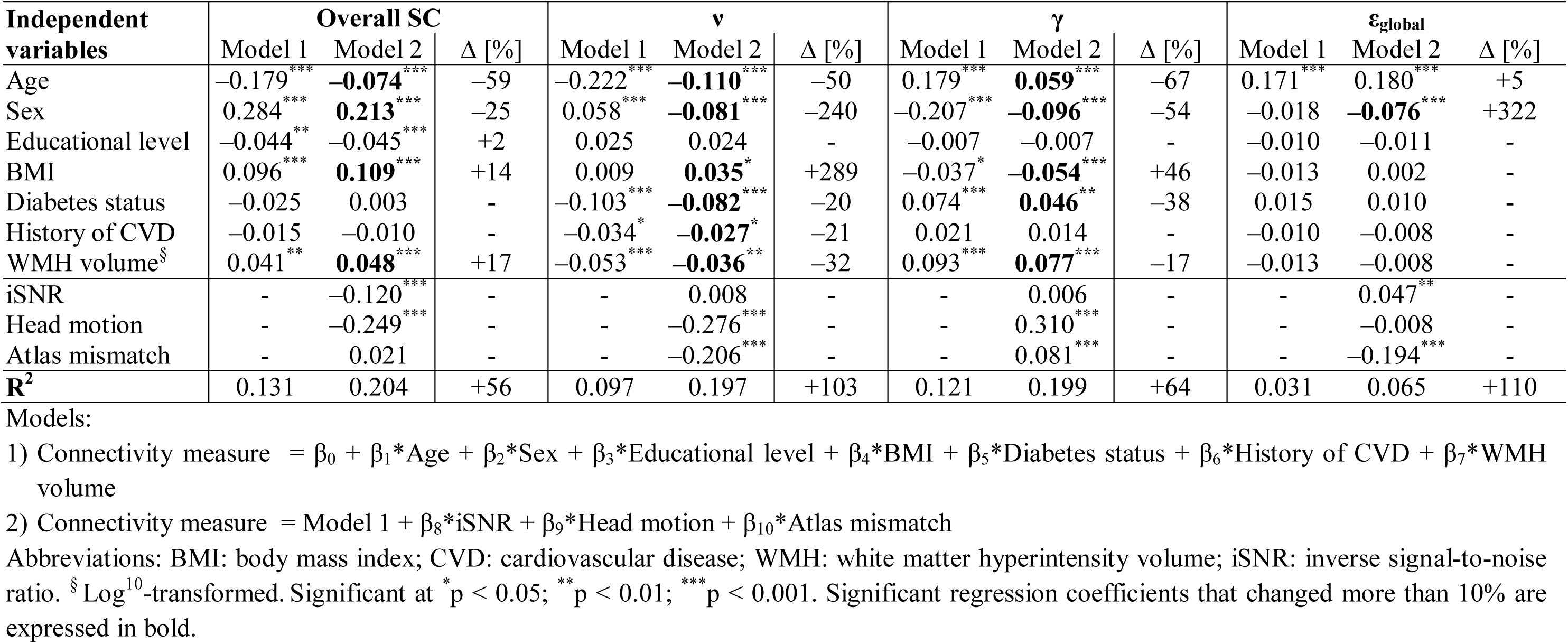
Standardized regression coefficients of the crude regression model between the structural connectivity measures at sparsity 0.80 and the demographic/clinical variables (Model 1), and the same model with additional adjustment for the diffusion MR image quality metrics (Model 2). For significant regression coefficients, also the percentage change is calculated.

**Table 2B.** Standardized regression coefficients of the crude regression model between the functional connectivity measures at sparsity 0.80 and the demographic/clinical variables (Model 1), and the same model with additional adjustment for the functional MR image quality metrics (Model 2). For significant regression coefficients, also the percentage change is calculated.

| Independent variables | Overall FC | | | $\nu$ | | | $\gamma$ | | | $\epsilon_{\text{global}}$ | | |
| --- | --- | --- | --- | --- | --- | --- | --- | --- | --- | --- | --- | --- |
| | Model 1 | Model 2 | $\Delta$ [%] | Model 1 | Model 2 | $\Delta$ [%] | Model 1 | Model 2 | $\Delta$ [%] | Model 1 | Model 2 | $\Delta$ [%] |
| Age | -0.003 | 0.015 | - | -0.129*** | -0.119*** | -8 | 0.050** | **0.042\*\*** | -16 | 0.063*** | **0.050\*\*** | -21 |
| Sex | -0.054** | **-0.062\*\*\*** | +15 | 0.006 | 0.003 | - | -0.006 | -0.012 | - | -0.113*** | **-0.095\*\*\*** | -16 |
| Educational level | 0.017 | 0.014 | - | 0.039** | 0.036* | -8 | -0.045** | -0.042** | -7 | 0.013 | 0.015 | - |
| BMI | -0.029 | -0.008 | - | -0.030* | **-0.005** | -83 | 0.001 | -0.017 | - | 0.094*** | **0.049\*\*** | -48 |
| Diabetes status | -0.037* | **-0.030** | - | -0.065*** | -0.060*** | -8 | 0.060*** | **0.054\*\*\*** | -10 | 0.013 | 0.008 | - |
| History of CVD | -0.016 | -0.017 | - | -0.040** | -0.041** | +3 | 0.009 | 0.010 | - | 0.010 | 0.011 | - |
| WMH volume<sup>§</sup> | -0.030* | **-0.027** | - | -0.058*** | -0.057*** | -2 | 0.021 | 0.018 | - | 0.000 | 0.002 | - |
| iSNR | - | -0.082*** | - | - | -0.052** | - | - | 0.075*** | - | - | 0.004 | - |
| Head motion | - | 0.048** | - | - | -0.010 | - | - | -0.035* | - | - | 0.127*** | - |
| Atlas mismatch | - | 0.027 | - | - | 0.004 | - | - | 0.016 | - | - | -0.029 | - |
| **R<sup>2</sup>** | 0.007 | 0.012 | +71 | 0.046 | 0.049 | +7 | 0.013 | 0.017 | +31 | 0.033 | 0.048 | +45 |

Models:
1) Connectivity measure = $\beta_0 + \beta_1 \cdot \text{Age} + \beta_2 \cdot \text{Sex} + \beta_3 \cdot \text{Educational level} + \beta_4 \cdot \text{BMI} + \beta_5 \cdot \text{Diabetes status} + \beta_6 \cdot \text{History of CVD} + \beta_7 \cdot \text{WMH volume}$
2) Connectivity measure = Model 1 + $\beta_8 \cdot \text{iSNR} + \beta_9 \cdot \text{Head motion} + \beta_{10} \cdot \text{Atlas mismatch}$
Abbreviations: BMI: body mass index; CVD: cardiovascular disease; WMH: white matter hyperintensity volume; iSNR: inverse signal-to-noise ratio. <sup>§</sup> Log<sup>10</sup>-transformed. Significant at \* $p < 0.05$ ; \*\* $p < 0.01$ ; \*\*\* $p < 0.001$ . Significant regression coefficients that changed more than 10% are expressed in bold.

Without adjustment for image quality, in particular higher age and male sex were associated with lower overall structural connectivity (β=–0.180, p<0.001; and β=0.284, p<0.001, respectively), lower average node degree (β=–0.222, p<0.001; and β=0.058, p<0.001, respectively), and higher clustering coefficient (β=0.179, p<0.001; and β=–0.207, p<0.001, respectively). With adjustment for diffusion MR image quality, the aforementioned associations with age and sex decreased by more than 26%. Without adjustment for image quality, higher age, but not sex, was associated with higher global efficiency (β=0.171, p<0.001; and β=0.018, p=0.204, respectively), while male sex was found to be associated with lower global efficiency (β=–0.076, p<0.001) after adjustment for image quality. Of note, significant associations were also observed for different combinations of demographic/clinical variables and the structural connectivity measures, with standardized regression coefficients generally <0.1 with relevant changes (>10%) after adjustment for image quality (See **Table 2A**).

Without adjustment for image quality, higher age and male sex were associated with lower functional average node degree (β=–0.129, p<0.001) and higher functional global efficiency (β=–0.113, p<0.001), respectively, and these associations did not change after adjustment for functional MR image quality. No other associations with |β|>0.1 were observed between the demographic/clinical variables and the functional connectivity measures (see **Table 2B**).

### Associations between quality and demographic variables

Standardized regression coefficients (β) between the demographic/clinical variables and each of the diffusion and functional MR image quality metrics, including the R^2^-value for the complete model, are listed in **Table 3A and 3B**. From the three diffusion MR image quality metrics (**Table 3A**), variance in head motion could be best explained by the demographic/clinical variables (R^2^=0.297) with age (β=0.368,p<0.001) and sex (β=–0.279, p<0.001), indicating more head motion at higher age and in men compared to women, as strongest covariates. Mismatch between diffusion MRI and brain atlas had the strongest association with sex (β=–0.303, p<0.001), indicating less mismatch in women compared to men. Conversely, the amount of variance in iSNR of the diffusion MRI that could be explained by the demographic/clinical variables was negligible (R^2^=0.033). From the three functional MR image quality metrics (**Table 3B**), variance in iSNR could be best explained by the demographic/clinical variables (R^2^=0.270) with BMI (β=0.408, p<0.001) as strongest covariate. Variance in head motion and atlas mismatch of the functional MRI were best explained by BMI (β=0.321, p<0.001) and sex (β=0.317, p<0.001), respectively. Scatterplots and histograms visualizing the strongest associations between quality metric and demographic variables are shown in **Figure 7**.

**Table 3A.** Standardized regression coefficients obtained from a linear regression model with forward selection between demographic/clinical variables and each of the quality metrics for the diffusion MRI.

| Independent variables | iSNR |  | Head motion |  | Atlas mismatch |  |
| --- | --- | --- | --- | --- | --- | --- |
| | $\beta$ | p-value | $\beta$ | p-value | $\beta$ | p-value |
| Age | **0.117** | <0.001 | **0.368** | <0.001 | **0.056** | <0.001 |
| Sex | **-0.071** | <0.001 | **-0.279** | <0.001 | **-0.303** | <0.001 |
| Educational level | -0.017 | 0.242 | 0.001 | 0.916 | -0.009 | 0.519 |
| BMI | **0.067** | <0.001 | **0.028** | 0.031 | **0.093** | <0.001 |
| Diabetes status | **0.032** | 0.042 | **0.095** | <0.001 | -0.023 | 0.129 |
| History of CVD | 0.004 | 0.774 | 0.020 | 0.093 | 0.009 | 0.514 |
| WMH volume<sup>§</sup> | **-0.038** | 0.009 | **0.048** | <0.001 | 0.017 | 0.224 |
| R2 | 0.033 |  | 0.297 |  | 0.113 |  |

Abbreviations: BMI: body mass index; CVD: cardiovascular disease; WMH: white matter hyperintensity volume; iSNR: inverse signal-to-noise ratio. <sup>§</sup> Log<sup>10</sup>-transformed. Significant regression coefficients ( $p < 0.05$ ) are expressed in bold.

**Table 3B.**
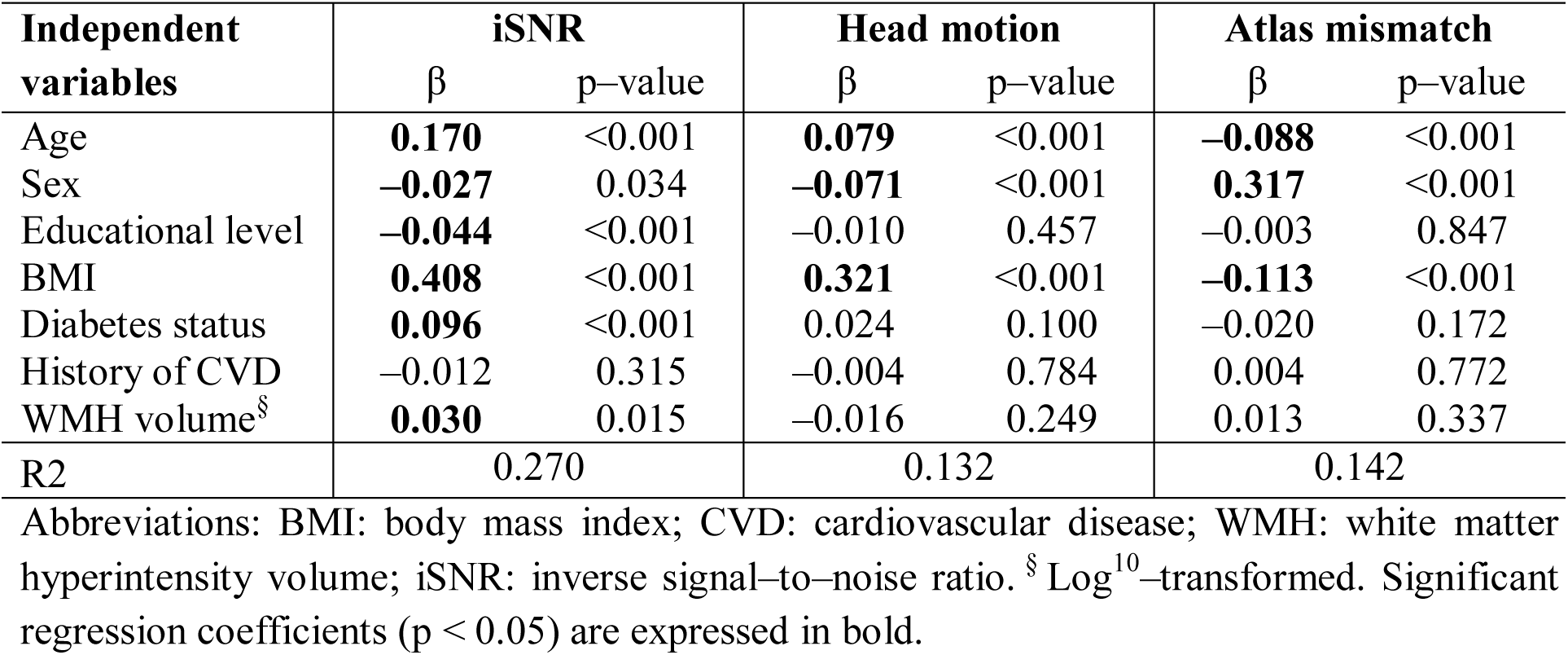
Standardized regression coefficients obtained from a linear regression model with forward selection between demographic/clinical variables and each of the quality metrics for the functional MRI.

**Figure 7.**
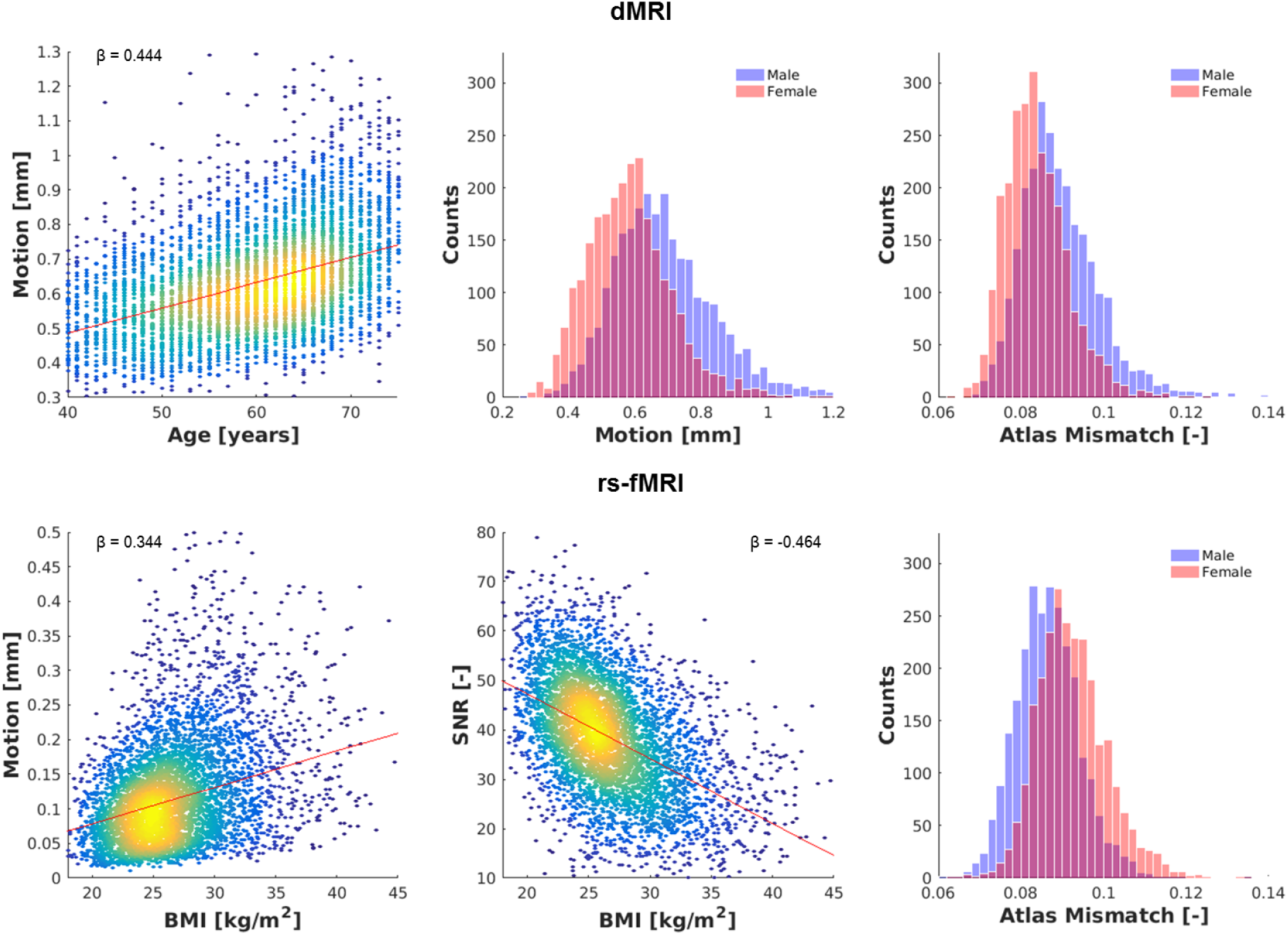
Examples of strongest determinants of quality metrics in the structural (top) and functional (bottom) connectivity pipeline. For structural quality, motion was best determined by age and sex, and atlas mismatch by sex. For functional quality, motion and iSNR were best determined by BMI, and atlas mismatch by sex. Note: for intuitiveness, SNR (expressed as – iSNR) is plotted instead of iSNR.

## Discussion

### Main findings

We extensively studied the association of dMRI and rs-fMRI quality with structural and functional connectivity measures, respectively, in 5,110 participants of The Maastricht Study. To summarize, we found a significant association between the dMRI and rs-fMRI quality metrics, i.e. in particular head motion and signal-to-noise ratio, respectively, and measures of structural and functional connectivity. Moreover, the image quality metrics affected the association between brain connectivity measures and demographic variables. Furthermore, our results showed that the image quality metrics were equally or even stronger determinants of brain structural and functional connectivity than demographic and/or clinical variables.

### Head motion

Head motion during the dMRI scan was most strongly associated with three of the four structural connectivity measures studied here, indicating it is an important potential confounder. To put the effect of head motion into perspective: every 0.1 millimetre of head motion during dMRI can be misinterpreted as a decrease in overall structural brain connectivity similar to 18.3 years of aging (see appendix for derivation).

For dMRI as well as rs-fMRI, the amount of head motion increased with age and was larger in men compared to women. These results are in line with current literature, as similar findings have been reported earlier (Geerligs et al., 2017; Huijbers et al., 2017; Savalia et al., 2017; Van Dijk et al., 2012). In addition, the amount of head motion during the rs-fMRI scan increased with BMI, which might be caused by the larger respiratory movements in persons with high BMI.

We also found that the amount of head motion was larger in the dMRI compared to the rs-fMRI scan (mean head motion 0.64 mm and 0.13 mm, respectively. An explanation for this finding might lie in the nature of the pulse sequences, for instance the longer echo and repetition times and the much stronger gradients of the dMRI scan, leading to notable table vibrations and head coil vibrations, which may amplify any distorting effects due to head motion, compared to the used rs-fMRI scan. Furthermore, the dMRI sequence was applied after the rs-fMRI sequence, at the end of the scan protocol. Hence, assuming that participants are more likely to move at longer scan times, this might explain the higher amount of head motion during dMRI compared to rs-fMRI.

Of note, the amount of head motion might not only be a confounder for the connectivity measures derived from the dMRI or rs-fMRI scans, but also for measures derived from structural MRI scans, i.e. T1w, FLAIR, SWI (Savalia et al., 2017). Thus, one might advise to adjust for image quality not only in statistical analyses on connectivity measures, but also in statistical analyses involving volumetric MRI measures.

### Brain atlas mismatch

Although the distribution of atlas mismatches in the studied population is highly comparable for dMRI and rs-fMRI (**Figures 4C and 4F**), atlas mismatch was associated with three out of four structural connectivity measures, but not with any of the functional connectivity measures. However, the dMRI-based structural connectivity measures rely on the geometric start and end (voxel) points as well as trajectories of streamlines connecting these points, which are likely more susceptible to geometric distortions than the rs-fMRI-based functional connectivity measures, which are based on spatially region-averaged signal time-series.

Since atlas mismatch and head motion are both associated with three out of four structural connectivity measures, they might have a common source, i.e. typical susceptibility artefacts due to the EPI sequences used during dMRI that are known to be highly prone to resonance offsets, e.g. magnetic susceptibility gradients, or B0 inhomogeneities (Bammer et al., 2001), for which ExploreDTI did not correct. The linear registration that we used to co-register the atlas to dMRI space, is only able to account for deformations caused by these susceptibility artefacts to a limited extent. Whether the use of a non-linear registration procedure, an individual-based atlas, or implementation of a more rigorous susceptibility correction method will lead to less mismatch, was beyond the scope of the current study.

### Signal-to-noise ratio (SNR)

SNR was weakly associated with one structural connectivity measure, i.e. overall SC, but with three out of four functional connectivity measures, indicating that, in addition to head motion, SNR could be a quality metric of interest in brain connectivity analyses. Interestingly, SNR in rs-fMRI, and to a lesser extent also in dMRI, decreased with BMI, which might have a physiological explanation as respiratory function is altered in obesity (Parameswaran et al., 2006), especially when scanned in the supine position, which may increase physiological-related noise (Kruger and Glover, 2001).

### Effect of image quality on associations between brain connectivity and demographic variables

Since the image quality metrics were significantly related to measures of structural and functional connectivity, it is apparent that they affect the associations between structural or functional connectivity and demographic/clinical variables. Indeed, without adjustment for image quality, the strength of the associations between structural connectivity and demographic variables, particularly age and sex and to a lesser extent BMI and WMH volume, differed by more than 25% compared to the model that adjusted for image quality. Interestingly, whereas the age- and sex-related associations with brain connectivity were weakened due to confounding effects of image quality, BMI-brain connectivity associations were actually strengthened when image quality was taken into account. A plausible explanation for this observation is currently still lacking. Although the ground truth structural connectivity in our study sample is unknown, the fact that age-, sex-, and BMI-related associations are affected by image quality underlines the importance of adjusting for it.

For functional connectivity associations with demographic/clinical variables, however, the changes obtained through adjustment for image quality were much smaller. An explanation for this discrepancy is that the gradients in dMRI are stronger than in rs-fMRI, hence any artefact is more pronounced in the dMRI and thus the effect of low image quality is stronger. Moreover, during the pre-processing of the rs-fMRI data, head motion is already taken into account by adding the motion parameters as nuisance regressors to the regression model when calculating temporal correlation between two brain regions.

### Validity structural and functional connectivity results

Analyses of the connectivity associations with demographic/clinical variables demonstrated that overall structural and functional connectivity, and hence node degree, decrease with age, whereas normalized clustering coefficient and global efficiency increase with age. This finding suggests that despite decreasing connectivity, whole brain network segregation as well as integration increases during aging. Decreased structural and functional connectivity during aging is consistent with the current consensus as summarized in a recent review (Damoiseaux, 2017). Less consensus, however, exists in the literature on the association of global efficiency with age. For example, structural and functional global efficiency were lower in older compared to young people (Achard and Bullmore, 2007; Zhao et al., 2015), or showed no difference between old and young people (Cao et al., 2014; Geerligs et al., 2015; Gong et al., 2009), whereas we found a slight positive association. The positive association between structural clustering coefficient and age that we found confirms the findings reported by Zhao et al. (Zhao et al., 2015). Yet, it has to be noted that the aforementioned findings have been reported in studies with relatively small sample sizes (n ≤ 126) compared to our study.

The validity of our structural and functional connectivity results is further supported by their dependency on sparsity. Both the structural and functional average node degree decreased with sparsity. This was as expected, since with increasing sparsity fewer connections are evaluated, and thus the number of possible connections to each node decreases as well. Structural and functional normalized clustering coefficient increased with sparsity. This effect, too, can be explained by the methodology used, because the proportion of connections between the nodes within its neighbourhood divided by the number of connections that theoretically could exist between them, will decrease with increasing sparsity. Conversely, the normalized clustering coefficient increases at increasing sparsity, because the clustering coefficient is normalized to a random network, for which the proportional decrease is larger.

In contrast, the normalized global efficiency over sparsity showed an opposite trend in the structural compared to the functional connectomes. This difference can be explained by varying number of intra- and interhemispheric connections taken into account in the structural and functional group-averaged connectomes over the sparsity range. While the percentage of interhemispheric connections in the functional group-averaged connectomes remains fairly constant (at 41-44%), this percentage decreases in the structural group-averaged connectomes from 25% at sparsity of 0.60 to 12% at sparsity of 0.90 (see **Supplemental Figure S5** in the appendix). As the connection-weights in the structural connectomes represent tract volumes, and since the structural connections taken into account are mostly short (intra-hemispheric) tracts with a small volume, the structural global efficiency is calculated using fairly low connection strengths that increases with sparsity, whereas the functional global efficiency is based on connections of high strength that increase with sparsity.

The result that fewer interhemispheric connections were taken into account in the structural compared to the functional connectomes, can be explained by the effect of length and shape of the tracts in whole-brain tractography (Jones, 2010). Since interhemispheric tracts are generally longer than intrahemispheric tracts, they are more difficult to track and are thus less likely to end up in the group-averaged connectome in favour of intrahemispheric connections.

### Strengths and limitations

Strengths of this study include the large number of participants and the acquisition of dMRI as well as rs-fMRI in these participants. However, there are also several limitations that are noteworthy to address. First of all, we could not implement any advanced correction for B0-field inhomogeneities or geometric distortions, e.g. FSL’s “topup” (Andersson et al., 2003), in the dMRI processing pipeline due to missing reversed phase-encode blips in the dMRI sequence. Hence, we were restricted to less advanced methods, such as (linear) registration to standard space. Consequently, the results relating to regions in the anterior frontal cortex and temporal lobe might therefore be less reliable as geometric distortions often occur in this location (Jezzard and Balaban, 1995). However, to our knowledge there is no reason to assume that these artefacts differ between subgroups, e.g. participants with and without T2DM, and therefore no bias has been introduced.

Second, we used an atlas template that is not participant-specific and as such may contribute to mismatch between the brain atlas and the participant’s dMRI or rs-fMRI. An individual-based brain parcellation, such as implemented in the FreeSurfer software (Destrieux et al., 2010), might result in better overlap with the participant’s dMRI or rs-fMRI. However, this requires substantially longer processing times, e.g. up to 20 hours per participant, as well as visual checks and manual intervention, whereas linear registration of the AAL2 atlas is robust and is typically completed within a minute.

## Conclusion

To conclude, we here describe the complete pipeline analyses for the assessment of the structural and functional brain connectivity in The Maastricht Study, including extensive quality assessment focused on the confounding effects of compromised image quality in population neuroimaging studies. Structural connectivity estimates were most strongly associated with head motion, while functional connectivity estimates were mainly influenced by signal-to-noise ratio. Moreover, image quality metrics had larger effects on brain connectivity estimates than demographic variables such as age or sex. Based on these findings, we recommend that statistical analyses of structural and functional brain connectivity and its associations with demographic or clinical variables should consider potential confounding effects of image quality.

## Acknowledgements

The authors are indebted to the participants for their willingness to participate in the study. In addition, the authors would like to thank Jos Slenter and Jan Jungerius from the Department of Radiology and Nuclear Medicine, Maastricht University Medical Centre, Maastricht, The Netherlands, for their assistance with source data retrieval, storage and processing.

This study was supported by the European Regional Development Fund via OP-Zuid, the Province of Limburg, the Dutch Ministry of Economic Affairs (grant 31O.041), Stichting De Weijerhorst (Maastricht, the Netherlands), the Pearl String Initiative Diabetes (Amsterdam, the Netherlands), CARIM, School for Cardiovascular Diseases (Maastricht, the Netherlands), School CAPHRI, Care and Public Health Research Institute (Maastricht, the Netherlands), NUTRIM, School of Nutrition and Translational Research in Metabolism (Maastricht, the Netherlands), Stichting Annadal (Maastricht, the Netherlands), Health Foundation Limburg (Maastricht, the Netherlands) and by unrestricted grants from Janssen-Cilag B.V. (Tilburg, the Netherlands), Novo Nordisk Farma B.V. (Alphen aan den Rijn, the Netherlands) and Sanofi-Aventis Netherlands B.V. (Gouda, the Netherlands).

## Appendices

## Appendix 1 Figure S1 Flow-chart of participants that were included in this study

**Supplemental Figure S1.**
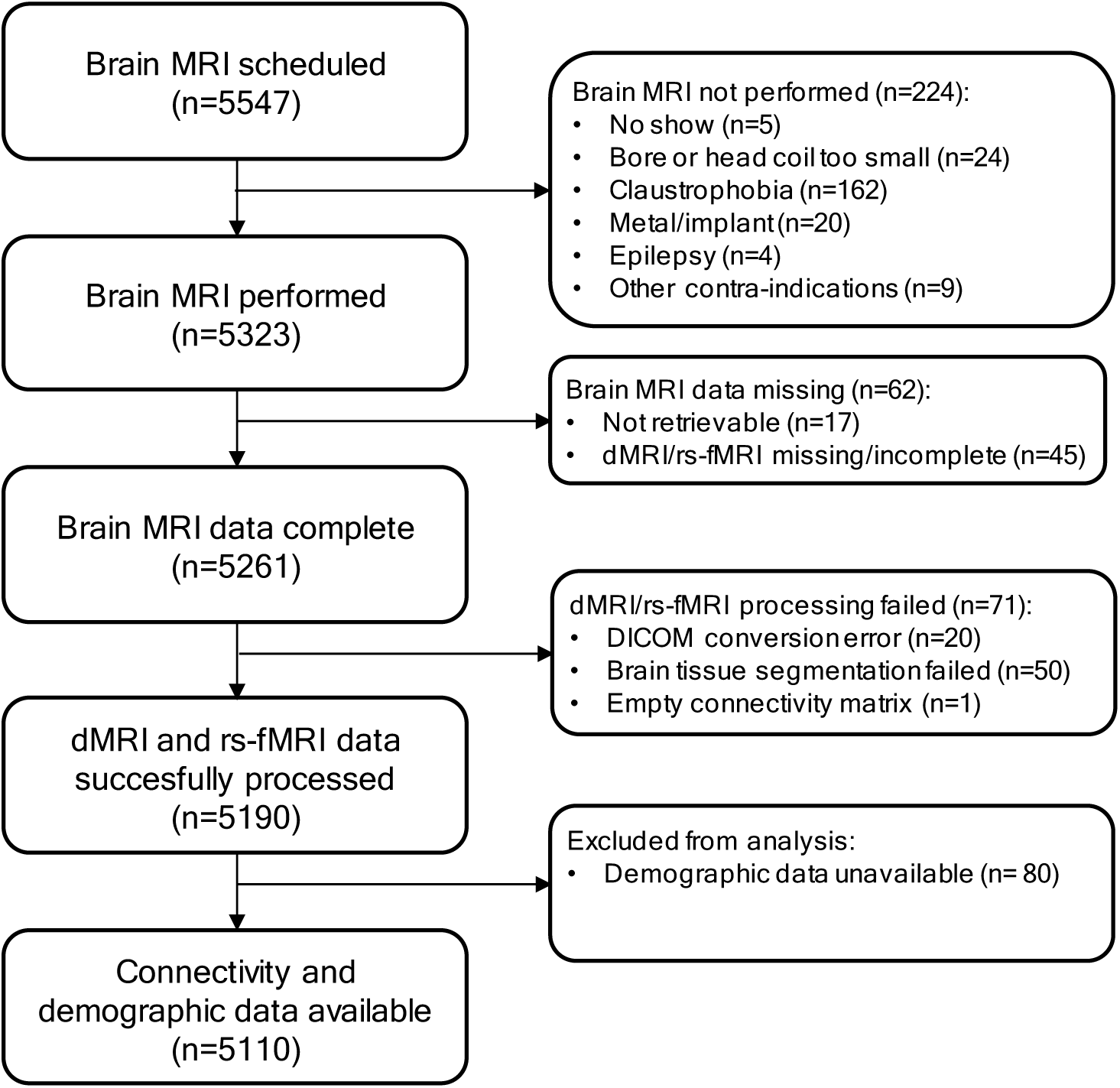
Flowchart showing the number of participants for whom both structural and functional brain connectivity measures could be calculated successfully, and demographic data were available.

## Appendix 2 Complete description of structural and functional connectivity processing pipeline

### Diffusion data pre-processing

dMRI data were anonymized and converted from DICOM to NIfTI format first using Chris Rorden’s dcm2nii tool (version 2MAY2016 64bit BSD License). After importing the NIfTI files into ExploreDTI v4.8.6 (PROVIDI lab, Image Sciences Institute, Utrecht, The Netherlands) (Leemans et al., 2009), eddy current and head motion correction was applied, while making sure the b-matrix was rotated accordingly (Farrell et al., 2007; Leemans and Jones, 2009). Next, white matter tracts were calculated using a constrained spherical deconvolution (CSD)-based deterministic tractography algorithm (Tax et al., 2014) at the following settings: 2 mm seed point resolution with seed points placed randomly throughout the whole brain; step size 1 mm; and maximum harmonic degree of 8 (Tournier et al., 2013). Stopping criteria were: fibre orientation distribution < 0.1; angle deviation > 30°; fibres leaving the brain mask; or fibre length < 50 mm or > 500 mm.

The automated anatomical labelling (AAL) atlas (Rolls et al., 2015), consisting of 94 (sub)cortical brain regions in the cerebrum, was linearly coregistered to the dMRI data using FLIRT (Jenkinson and Smith, 2001) in FMRIB Software Library (FSL) 5.0.10 (FMRIB Analysis Group, University of Oxford, Oxford, U.K.) and imported into ExploreDTI.

Subsequently, for each pair of brain regions, the connection strength was defined as the tract volume (number of voxels visited by a tract multiplied by the voxel size) divided by the ICV if two or more tracts were found between the two brain regions, otherwise the connection was considered as absent and the connection strength set to zero (Vaessen et al., 2010). Also the diagonal elements (i.e. self/self-connections) were set to zero. Finally, this resulted in a symmetric 94×94 connectivity matrix, i.e. the participant’s structural connectome (SC), were each row and column represents a brain region and each element represents the relative tract volume between two regions.

### Functional data pre-processing

Rs-fMRI data were anonymized and converted from DICOM to NIfTI format using Chris Rorden’s dcm2nii tool (version 2MAY2016 64bit BSD License). To account for magnetization stabilization, the first ten seconds of data (equivalent to the first five volumes) were removed, and the remaining rs-fMRI volumes were corrected for field inhomogeneities using FSL 5.0.10 (FMRIB Analysis Group, University of Oxford, Oxford, U.K.) (Zhang et al., 2001). The rs-fMRI data were then imported into Statistical Parametric Mapping (SPM) 12 (The Wellcome Trust College London, London, U.K.). Because the rs-fMRI data were acquired in an interleaved spatial order, slice-timing correction (with the second slice as reference since this slice was acquired first) was applied before head motion correction (Soares et al., 2016). To improve the signal-to-noise ratio, the rs-fMRI images were spatially smoothed using a Gaussian kernel (full width at half maximum = 8 mm). Last, the rs-fMRI data were temporally filtered using a FSL’s band-pass filter (0.01 to 0.1 Hz) (Biswal et al., 1995) to remove possible respiratory and signal drift effects and to focus on the spontaneous low-frequency fluctuations.

The participant’s structural T1w images, including the masks of the CSF and WM, as well as the AAL atlas (Rolls et al., 2015) were linearly coregistered to the rs-fMRI data using FSL’s FLIRT (Jenkinson and Smith, 2001), and an averaged time-series in each brain region as well as in the CSF and WM was calculated from the per-voxel time-series in each region.

Subsequently, for each pair of brain regions, a Pearson’s correlation coefficient was calculated using linear regression of the averaged time-series of each region, with the averaged time-series in the CSF and WM and the motion correction parameters as nuisance regressors in MATLAB Release 2016a (The Mathworks Inc., Natick, Massachusetts, U.S.). This resulted in a symmetric 94×94 correlation matrix, i.e. the participant’s functional connectome (FC). In the FC, each row and column represent a brain region and each element represents the temporal correlation between two regions. Last, the diagonal elements (self/self-connections) and negative correlations, which are considered not representing any meaningful connections, were set to zero (Smith et al., 2011), resulting in a correlation weighted, undirected network.

### Hardware and software

All structural and functional connectivity analyses were performed on a dedicated computer cluster containing four nodes with each a Intel Xeon E3-1245v3 3.40 GHz 8-core processor, 32 GB of DDR4 RAM and a 250 GB SSD.

Processed data were stored on dedicated storage servers equipped with in total eight 10 TB SATA 6.0 Gb/s hard disks. To obtain a high level of data safety, the disks were configured in two RAID 6 arrays, thus giving protection against simultaneous failure of two disks and resulting in 20 TB of usable disk space.

Each node was loaded with an image that contained the operating system, drivers and dedicated software. The operating system was 64-bit Scientific Linux release 6.8 (Carbon) based on Linux kernel 2.6.32-642.11.1.el6.x86_64 and GNOME 2.28.2. The software included in the image is listed in **Table S1**.

**Table S1.** Overview of neuro-imaging software used in the structural and functional connectivity processing pipelines.

| Software package | Version | Release date | RRID |
| --- | --- | --- | --- |
| dcm2nii | 2MAY2016 | 02/05/2016 | SCR_014099 |
| Matlab including the Image Processing Toolbox | 2016a | 01/01/2016 | SCR_001622 |
| Brain Connectivity Toolbox | 2017_01_15 | 15/01/2017 | SCR_004841 |
| ExploreDTI | 4.8.6 | 23/02/2017 | SCR_001643 |
| FMRIB Software Library | 5.0.10 | 25/04/2017 | SCR_002823 |
| Statistical Parametric Mapping | 12 rev 6906 | 20/10/2016 | SCR_007037 |

RRID: research resource identifier as reported by https://scicrunch.org

### Processing times and size of generated data

Average processing times per participant in the structural and functional network connectivity pipeline were 1h33m and 0h50m, respectively. With the hard- and software used in this study, it took approximately 9 days to process the dMRI and rs-fMRI DICOM data of all participants. The amount of data generated per participant in the structural and functional network connectivity analysis were 847 MB and 744 MB, respectively, yielding a total of approximately 3.6 TB of generated data for the complete structural and functional connectivity analyses.

## Appendix 3 Table S3A and S3B Standardized regression coefficients and goodness of fit parameters between the connectivity measures and the image quality metrics

**Supplemental Table S3A.**
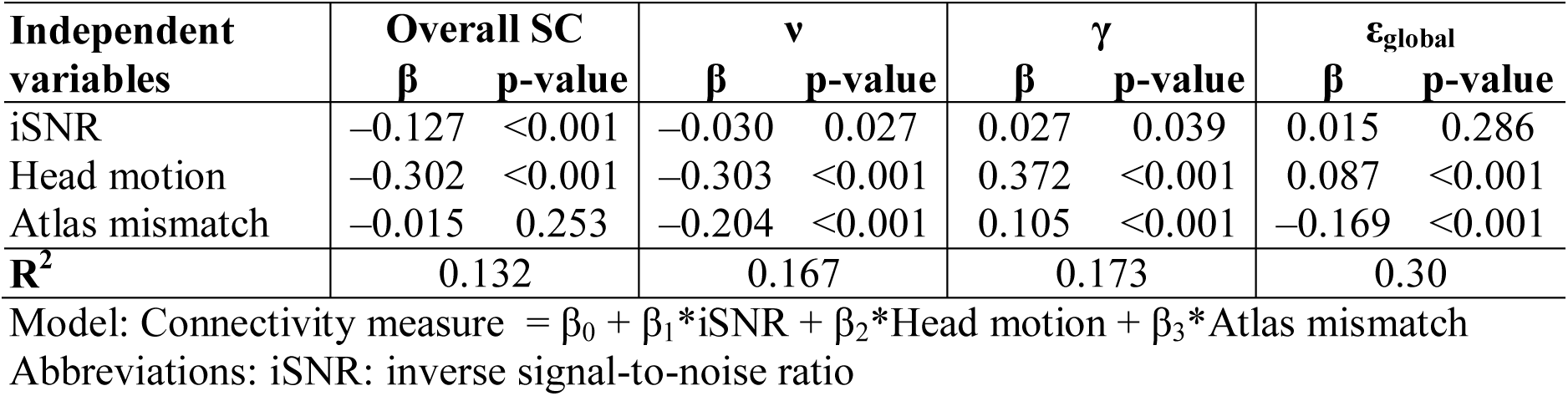
Standardized regression coefficients and goodness of fit parameters between the structural connectivity measures and the image quality metrics of the diffusion MRI.

**Supplemental Table S3B.** Standardized regression coefficients and goodness of fit parameters between the functional connectivity measures and the image quality metrics of the functional MRI.

| Independent variables | Overall FC | | $\nu$ | | $\gamma$ | | $\epsilon_{\text{global}}$ | |
| --- | --- | --- | --- | --- | --- | --- | --- | --- |
| | $\beta$ | p-value | $\beta$ | p-value | $\beta$ | p-value | $\beta$ | p-value |
| iSNR | -0.093 | <0.001 | -0.114 | <0.001 | 0.105 | <0.001 | 0.043 | 0.008 |
| Head motion | 0.053 | 0.001 | -0.022 | 0.176 | -0.028 | 0.087 | 0.127 | <0.001 |
| Atlas mismatch | 0.011 | 0.445 | 0.020 | 0.163 | 0.011 | 0.451 | -0.075 | <0.001 |
| $R^2$ | 0.006 | | 0.017 | | 0.008 | | 0.032 | |

Model: Connectivity measure = $\beta_0 + \beta_1 \cdot \text{iSNR} + \beta_2 \cdot \text{Head motion} + \beta_3 \cdot \text{Atlas mismatch}$
Abbreviations: iSNR: inverse signal-to-noise ratio

## Appendix 4 Explanation 0.1 mm of head motion equivalent to 18.3 years of aging

From the standard deviations for age and head motion in dMRI as reported in Table 1 and the standardized regression coefficients as reported in Table 2A, we can calculate how many years of aging has the equivalent effect on overall structural connectivity as a given amount of millimetres head motion.

From Table 2A we get the standardized regression coefficients for the complete model:

Overall structural connectivity = β_0_ **– 0.074*Age** + 0.213*Sex – 0.045*Educational level + 0.109*BMI + 0.003*Diabetes status – 0.010*History of CVD + 0.048*WMH volume – 0.120*iSNR **– 0.249*Head motion** + 0.021*Atlas mismatch

This can be interpreted as *“for one SD years increase in age, the overall structural connectivity will decrease by **0.074 SD**, and for one SD mm increase of head motion, the overall structural connectivity will decrease by **0.249 SD**”*.

From Table 1 we get the standard deviations for age and head motion, which are 8.7 years and 0.16 mm, respectively. Thus, for each **8.7 years** of aging, overall structural connectivity decreases by 0.074 SD, and for each additional **0.16 mm** of head motion, overall structural connectivity decreases by 0.249 SD.

To have a decrease of one SD in overall structural connectivity, we would need 117.6 (= 8.7 years*1SD/0.074SD) years of aging, or 0.643 (= 0.16 mm*1SD/0.249SD) millimetres of head motion.

And thus 117.6 years of aging has the equivalent effect on overall structural connectivity 0.643 mm of head motion, which is the same as: 18.3 (117.6/6.43) years of aging has the equivalent effect on overall structural connectivity 0.1 (= 0.643/6.43) mm of head motion.

## Appendix 5 Figure S5 Structural and functional group-averaged connectivity matrices at different sparsities

**Supplemental Figure S5.**
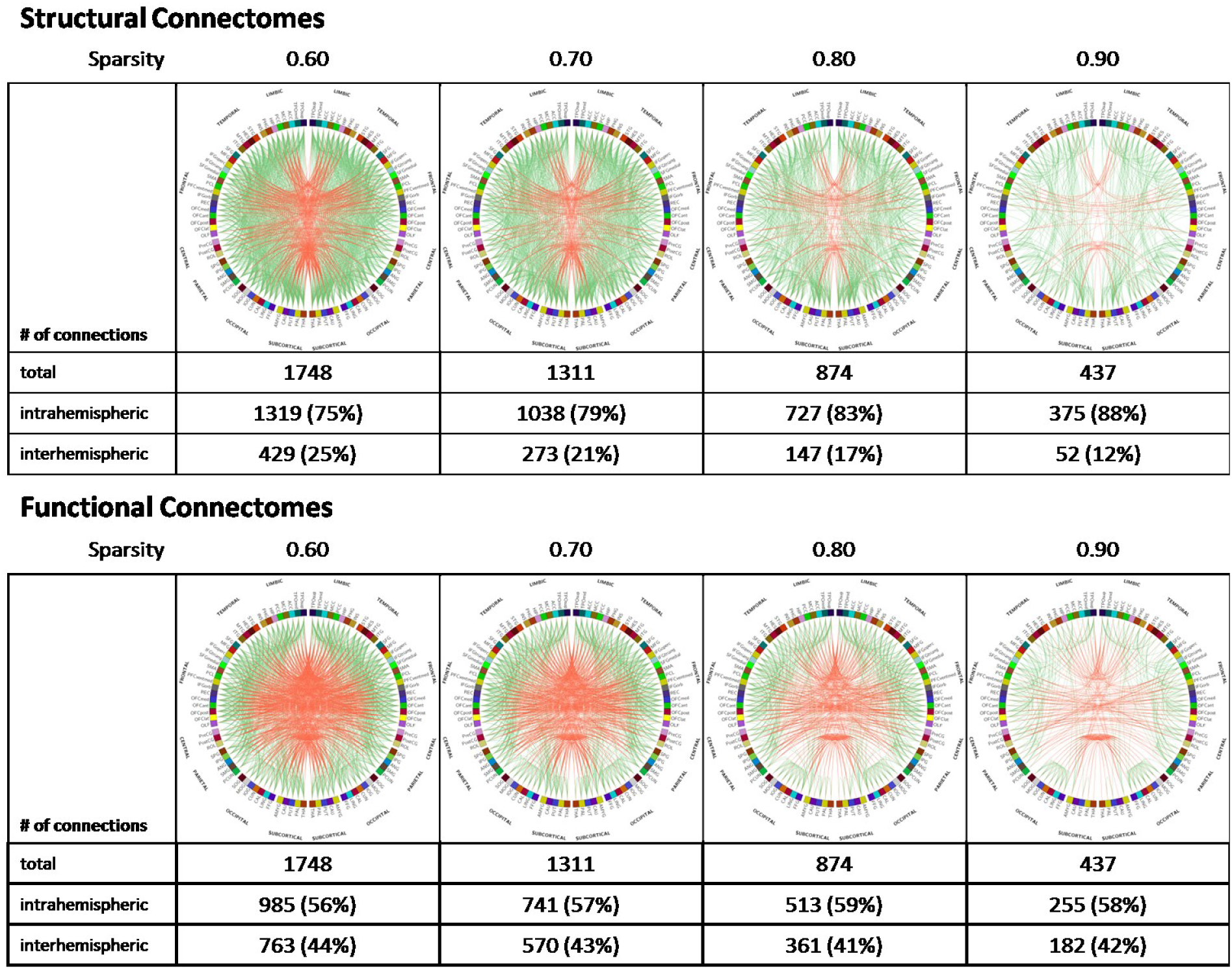
Structural (top) and functional (bottom) group-averaged connectomes at sparsity 0.60 to 0.90. Whereas the ratio of intra- and interhemispheric connections remains fairly stable for the functional connectomes, this ratio increases rapidly for the structural connectomes.

